# Non-B DNA-Informed Mutation Burden as a Marker of Treatment Response and Outcome in Cancer

**DOI:** 10.1101/2024.01.04.574248

**Authors:** Qi Xu, Jeanne Kowalski

## Abstract

**Background:** Genomic instability plays a key role in tumorigenesis and cancer research, with Tumor Mutation Burden (TMB) being a crucial biomarker quantifying total mutation to indicate therapeutic effectiveness, particularly in immunotherapy. However, TMB is not always a reliable predictor of treatment response and displays heterogeneity. Non-B DNA, alternative DNA forms have the potential to increase susceptibility to mutations that lead to the development of cancer. The tendency of these structures to induce mutations highlights their critical role in cancer onset and advancement, indicating their potential merit when combined with mutation information for enhanced markers in cancer with potential novel insights.

**Methods and findings:** We introduce two novel markers, “nbTMB” (non-B-informed tumor mutation burden) and “mlTNB” (mutation-localized-informed tumor non-B burden). We show in three separate case studies applying these markers the following findings: 1) nbTMB informs on survival heterogeneity among TMB-High patients undergoing immunotherapy whereas TMB is unable to further differentiate; 2) nbTMB informs on altered cisplatin sensitivity among ovarian cancer patient-derived cell lines whereas TMB is unable to differentiate; and 3) mlTNB informs on survival heterogeneity among early stage pancreatic cancer progressors in whom other markers of genomic instability fail to differentiate.

**Conclusions:** These novel markers offer a nuanced approach in which to enhance our current understanding of treatment responses and outcomes in cancer, underscoring the need for a more comprehensive exploration of the interplay between non-B and B-DNA features.

## Introduction

Genomic instability, marked by numerous genetic changes due to impaired DNA repair processes, stands as a hallmark of cancer (Andor et al. 2017). It plays an important role in contributing to the cancer heterogeneity, therapeutic resistance, and association with poor prognosis (Negrini et al. 2010; Shen 2011). Tumor Mutation Burden (TMB) stands as a crucial metric in oncology, quantifying the total mutations within tumor. It serves as a biomarker for the effectiveness of immunotherapies, particularly immune checkpoint inhibitors(Valero et al. 2021). A higher TMB often indicates a more favorable response to immunotherapies, providing a vital tool for clinicians in customizing therapeutic strategies and anticipating treatment outcomes (Karamitopoulou et al. 2022). However, the use of TMB and its threshold is not uniform across different cancer types, indicating the need for more nuanced exploration.

Within the genomic landscape, non-B DNA structures emerge as notable entities (Wang and Vasquez 2023; Xu and Kowalski 2023). These structures deviate from the conventional B-DNA double helix to form alternative structures (Wang and Vasquez 2023). They disrupt the standard processes of DNA replication and transcription, thereby laying the foundation for genetic instability. The propensity of these structures to induce mutations underscores their potentially critical role in cancer initiation and progression (Georgakopoulos-Soares et al. 2018; Bacolla et al. 2019; Guiblet et al. 2021; McGinty and Sunyaev 2023). The propensity of these structures to induce mutations underscores their critical role in cancer initiation and progression (Georgakopoulos-Soares et al. 2018; Bacolla et al. 2019; Guiblet et al. 2021; McGinty and Sunyaev 2023). Their increased susceptibility to change gives rise to an abundance of population variants linked to non-B DNA motifs and an amplified frequency of somatic mutations at these sites, notably in cancer contexts. Even though numerous variants tied to non-B DNA motifs may not have a profound impact, these motifs play a pivotal role in the genetic diversity of the human genome (Bansal et al. 2022; Makova and Weissensteiner 2023). As a result, they stand out as primary areas of interest for disease development and genetic discrepancies (Bansal et al. 2022). It is essential to factor in the importance of non-B DNA motifs for predicting mutation frequencies and evaluating potential disease risks, while developing new biomarkers in the context of cancer.

Our investigation unveils two novel biomarkers: nbTMB (non-B-informed tumor mutation burden) and mlTNB (mutation-localized total non-B DNA burden), aiming to quantify the multi-dimensions of genomic instability using both tumor mutations and non-B DNA at a sample level. First, nbTMB, non-B informed TMB, quantifies mutations co-localized non-B forming regions (**Figure. 1a**). We further calculate nbTMB percentage (nbTMBp) to describe the proportion of tumor mutations co-localized with non-B DNA motifs, relative to (total) TMB. Second, mlTNB refers to mutation-localized total non-B DNA burden as a quantification of non-B DNAs. Different from nbTMB (mutation quantification), mlTNB, as the non-B marker, evaluates the counts of non-B motifs containing mutation sites with its regions (**Figure. 2a**).

**Figure. 1.**
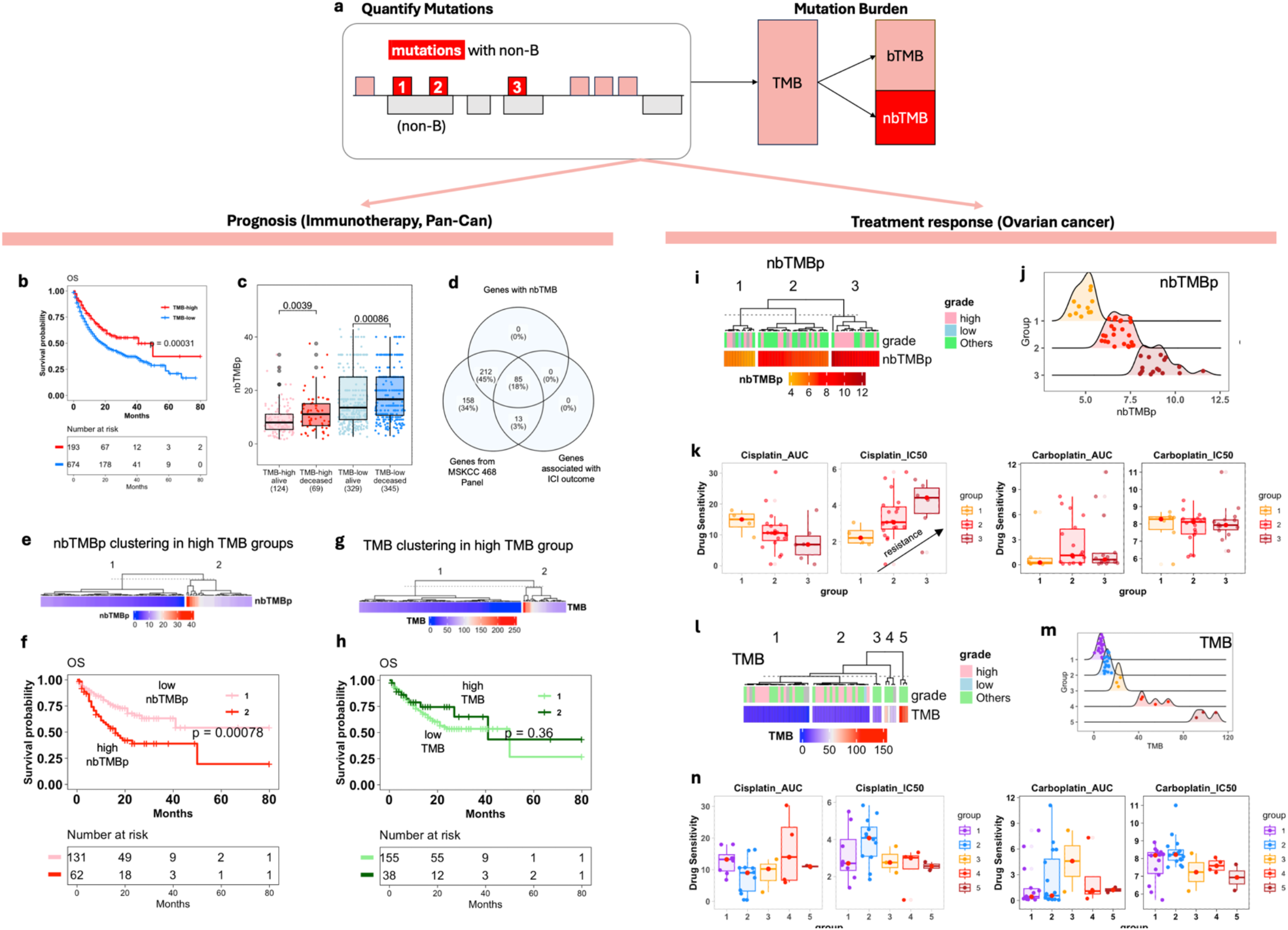
nbTMBp predicts patient outcome and drug sensitivities. **(a)** Overview of non-B-informed mutations quantification and use of nbTMBp in TMB decomposition. **(b)** Kaplan-Meier survival curve demonstrates that patients with elevated TMB have significantly improved overall survival compared to those with lower TMB when subjected to immunotherapy. **(c)** Among pan-can patients categorized by TMB levels (high or low), a further distinction into “alive” and “deceased” based on overall survival (OS) reveals that the deceased cohort consistently exhibits a higher nbTMBp percentage across both TMB categories. **(d)** The Venn diagram illustrates the overlapping genes between the MSKCC-Panel-468, those with non-B localized mutations, and the genes associated with the ICI-outcome (immune checkpoint inhibitors) signature. **(e-f)** In the TMB-high patient cohort, individuals with elevated nbTMBp exhibit reduced survival rates relative to those with lower nbTMBp. **(g-h)** Further categorization of the TMB-high patient group by their TMB levels (high or low) reveals no significant difference in survival outcomes. **(i-j**) Cell line clustering based on nbTMBp shows three distinct clusters, each characterized by varying levels of nbTMBp. **(k**) nbTMBp shows a linear trend of increasing drug resistance of Cisplatin. This is evident in both IC50 metrics (where a lower value indicates increased sensitivity) and dose-response AUC (where a higher value indicates increased sensitivity). Such a correlation is absent in the case of another platinum-based compound Carboplatin. **(l-n)** TMB alone does not show correlation with drug sensitivities of Cisplatin and Carboplatin in ovarian cancer cell lines.

**Figure. 2.**
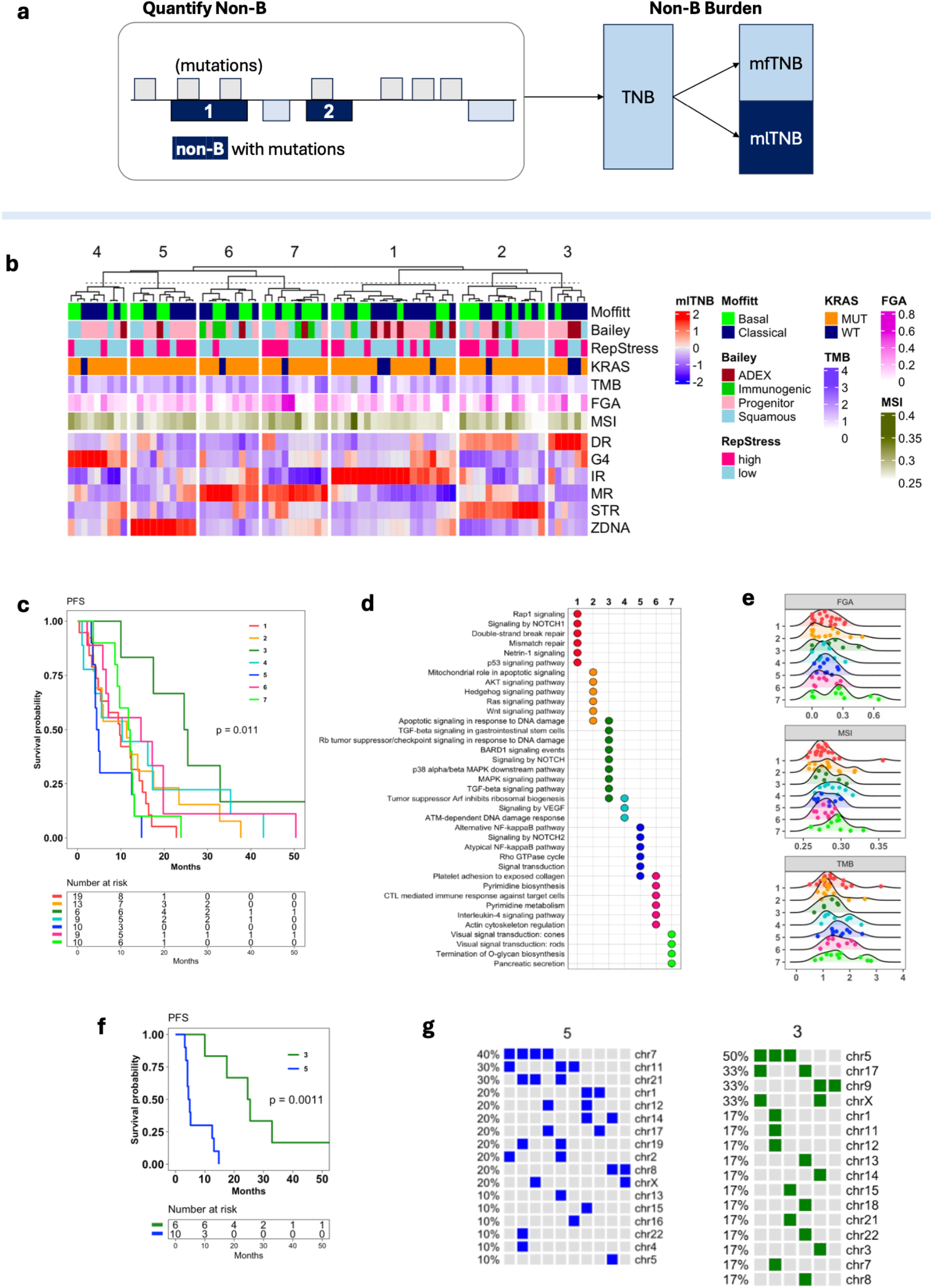
mlTNB predicts prognosis in early stage pancreatic cancer. **(a)** Schematic representation of mutation-localized total non-B burden categorization: TNB, mlTNB (mutation-localized), and mfTNB (mutation-free). **(b)** Clustering analysis of early-stage pancreatic cancer patients with progression, resulting in seven distinct clusters characterized by varying mlTNB burdens from different non-B types: direct repeats (DR), G-quadruplexes (G4), inverted repeats (IR), mirror repeats (MR), short tandem repeats (STR) and Z-DNA (ZDNA). **(c)**Kaplan-Meier progression-free survival (PFS) analysis for the identified seven patient clusters. **(d)** Pathway enrichment analysis highlighting key gene mutation signatures across the clusters. **(e)**The distribution of FGA, TMB and MSI (MANTIS score). All three metric did not show significant difference across seven groups. These sample have low TMB (median TMB < 2), low FGA (median < 30%), and are MSS (median MANTIS score < 0.3) **(f)** Comparative PFS survival curves for the cluster with the longest median PFS (mlTNB-DR, cluster 3) and the one with the shortest median PFS (mlTNB-ZDNA, cluster 5). **(g)** Chromosomal distribution of non-B mutation co-localizations that contribute to mlTNB burden.

We explored the heterogeneity within TMB-High patients by using nbTMB to investigate its role as a marker associated with post-immunotherapy survival. Our results lend support for the potential use of nbTMB to differentiate immunotherapy patient outcomes and thus inform on patients who might otherwise show unfavorable response despite their tumor having a high TMB. We next explored the use of nbTMB as a marker of altered cisplatin drug sensitivity in ovarian cancer. Our findings showed support for the further exploration of nbTMB as a marker that warrants further study for its potential in defining a molecular selected ovarian patient population of potential cisplatin treatment benefit. Lastly, we explored the use of mlTNB to quantify non-B burden and examine its association with survival in early-stage pancreatic cancer. Our results lend support to the further study of mlTNB as a differentiating marker of survival that may be used for pancreatic cancer patient outcome risk stratification.

## Results

### The design of nbTMB and mlTNB, based on non-B and mutation co-localization

Our investigation unveils two novel markers, nbTMB and mlTNB, aiming to quantify the multi-dimensions of non-B DNA motifs and the co-localized mutation sites. We intended to demonstrate their ability to act as localized mutation markers and gauge their utility as biomarkers across various cancer types.

The two markers are calculated for each tumor profile at sample level. The mutation signatures are extracted from each tumor profile. The genomic-wide non-B forming region are further overlapped with the mutated regions for each tumor profile. Using the overlapped region by counting separately the number of mutation and non-B motifs involved, we are able to derive the two metrics, nbTMB and mlTNB, as the new biomarker to reflect the interplay between mutation and non-B DNA. The metric was further refined by optional normalizations to predict patient prognosis and treatment responses.

nbTMB quantifies mutations within the realm of non-B as a non-B informed tumor mutation burden (**Figure. 1a**). The mutation signatures are extracted from each tumor profile. The genomic-wide non-B forming region is further overlapped with the mutated regions for each tumor profile. Further, we calculate nbTMB percentage (nbTMBp) to describe the proportion of tumor mutations co-localized with non-B DNA structures, relative to total tumor mutations.

On the other hand, mlTNB refers mutation localized non-B DNA burden as a quantification of non-B DNA motifs. Different from nbTMB, the marker, mlTNB, focuses on the counts of non-B motifs that contain mutation sites (**Figure. 2a**). Due the various non-B types, the mlTNB is further calculated by non-B motif types. For the comparison across non-B types and across samples, the burden value will also be normalized by both the number of mutations and the motif library size.

### nbTMB differentiates survival among TMB-High patient post-immunotherapy

We first describe a Pan-Can immunotherapy analyses in which nbTMBp appears linked with prognosis. High TMB has been reported to be associated with improved immunotherapy response (Samstein et al. 2019; Fusco et al. 2021; Rousseau et al. 2021). However, within TMB-high patients, outcomes remain heterogenous. Herein, we explore the heterogeneity with TMB high/low groups using nbTMBp to investigate its role as a biomarker associated with post-immunotherapy survival.

We analyzed the mutation data from patients who underwent immunotherapy based on the MSK-IMPACT study from 11 different cancer types (Samstein et al. 2019). Within each caner type, using the 80^th^ percentile of TMB, we assigned patients into TMB -high and -low groups and compared their overall survival (OS) (**Figure. 1b**). We further stratified patients within each of TMB-High and -Low groups based on their status – alive/censored or deceased. We defined nbTMB for each patient sample by quantifying the numbers of mutations co-localized with non-B forming regions (Cer et al. 2013). When comparing median nbTMBp across groups, the TMB-high group had a lower nbTMBp overall, relative to the TMB-low group (**Figure. 1c**). Within each TMB classified group, median nbTMBp was significantly higher in deceased patients, irrespective of their high/low status (**Figure. 1c**). A gene-level analysis of immune response signatures (Long et al. 2022) revealed an 86% overlap between mutations co-localized with non-B motifs and immune checkpoint inhibitor-outcome-linked genes (n=98) (**Figure. 1d**).

Next, we performed clustering on nbTMBp within the TMB-High patients, which revealed two patient subgroups (**Figure. 1e**), one with median nbTMPp of around 10% that was associated with significantly (p < 0.01) shorter OS as compared to TMB-High patients with less than 10% nbTMBp (**Figure. 1f**). For comparison, the same analysis was applied to TMB in which no significant OS difference was observed (**Figure 3g-h**). Altogether, our findings lend support for the further study of nbTMBp as a potential marker of differential survival within TMB-high patients on immunotherapy. These results may reflect the potential contribution from non-B DNA genomic instability in some patients that have poor survival, despite having high TMB.

### Increasing nbTMB is associated with decreased cisplatin sensitivity in ovarian cancer

We next explore the use of nbTMBp as a marker of altered cisplatin drug sensitivity in ovarian cancer. Cisplatin resistance is a major hurdle in effectively treating ovarian cancer (Mansouri et al. 2003). Although cisplatin is commonly used for ovarian cancer treatment, drug resistance often arises due to a faulty apoptotic process, reducing treatment effectiveness (Parker et al. 1991; Herr and Debatin 2001; Makin and Dive 2001; Song et al. 2022; Havasi et al. 2023). Among the 57 ovarian cell lines with mutation profiles(Ghandi et al. 2019), ∼40% have TMB greater than ten and a median fraction of genome altered (FGA) of ∼50%, which supports the potential role of genomic instability in treatment resistance. Investigating how cells signal in response to chemotherapy from the non-B DNA perspective of genomic instability may shed light on treatment outcomes.

We defined an ovarian cell line specific mutation signature and corresponding nbTMBp for use in a cluster analysis that identified three cell line groups of varying (low to high) nbTMBp (**Figure. 1i-j**). Median nbTMBp significantly differed among the three clustered cell line groups. Tests of association between TMB, FGA and tumor grade with nbTMBp-derived cell line clusters lacked significance, as did a correlation between TMB and nbTMBp among the ovarian cell lines. We examined the effect of clusters on cisplatin drug sensitivity in which increasing nbTMBp was significantly associated with decreasing cisplatin sensitivity. This finding was consistent for dose-response AUC with cisplatin (**Figure. 1k**). For comparison, we performed the same analyses on carboplatin sensitivity which did not show the same result, suggesting a cisplatin specific nbTMBp effect. Additionally, the use of TMB in a cluster analysis (**Figure. 1l-m**) failed to show a similar result (**Figure. 1m**). Altogether, our findings show support for the further exploration of nbTMBp as a potential marker of cisplatin sensitivity that may help to explain resistance when all other markers indicate otherwise.

### mlTNB differentiates survival among early-stage pancreatic patient progressors

In contrast to nbTMBp, we next explore the use of mlTNB to quantify non-B burden and its association with survival in pancreatic adenocarcinoma (PAAD). PAAD is a highly aggressive cancer with poor outcome. Existing genomic instability measures have not proven informative in differentiating survival into clinically translatable patient groups for risk stratification (Raphael et al. 2017). As opposed to focusing on mutation numbers, mlTNB quantifies non-B DNA regions that contain mutation sites which can be classified according to non-B structure types to provide a more nuanced perspective (Xu and Kowalski 2023) (**Figure. 2a**).

Using the mutation profiles of 76 TCGA early-stage pancreatic patients who progressed, we quantified mlTNB for each sample and used it in a cluster analysis resulting in seven patient groups with differentiated non-B structure types (**Figure. 2b**) that significantly differed in progression-free survival (PFS) (**Figure. 2c**). Patients with high mlTNB characterized mainly by direct repeat regions (high mlTNB-DR burden) was associated with the longest PFS (n = 7, median = 25 months), while patients with high mlTNB in Z-DNA regions had the shortest (n = 10, median = 5 months). PFS among other groups were similar: PFS among other groups were similar: the patient group with high mlTNB from short tandem repeat regions (STR) (n = 13, median = 11 months); the sample group with mlTNB from mirror repeats (MR) without inverted repeats (IR) (n = 7, median = 12 months); and the group with MR with IR (n = 9, median = 15 months).

The high mlTNB-DR burden patients had mutation signatures enriched in MAPK and Notch signaling pathways, as compared to the other clusters enriched with double-stranded break and mismatch repair (cluster 1, IR), hedgehog and WNT signaling (cluster 2, STR), and interleukin-4 signaling (cluster 6, MR) pathways (**Figure. 2d**). There was a lack of significant association between mlTNB clusters with age, race, sex, PAAD subtypes (Moffitt et al. 2015; Bailey et al. 2016), KRAS mutation status, and tumor purity. Additionally, there was no significant association between the non-B DNA clusters with markers of genomic instability, TMB, FGA, and MSI-score (measured by MANIS score) (Kautto et al. 2017), whose distributions were similar across clusters and on average, were low in value (**Figure. 2e**). Specifically, the median TMB was 1.43, the median FGA, 0.13, and the median MSI-score was 0.28 across all groups. In the shortest PFS (high mlTNB-ZDNA, cluster 5) (**Figure. 2f**), chromosome 7 had the highest prevalence of non-B mutation co-localization, while the longest PFS (high mlTNB-DR, cluster 3) patient group non-B mutation co-localization resided mainly on chromosome 5 (**Figure. 2g**). Our results lend support to the further study of mlTNB as a differentiating marker of survival in PAAD patients that may further inform on their heterogeneous response to treatment.

## Discussion

We explored the integration of non-B DNA information within mutation burden in cancer by the introduction of two makers: nbTMB and mlTNB. Our results show the added value from inclusion of non-B DNA data, lending support to the multi-dimension roles of genomic instability arising from non-B structures and their potential impact on cancer prognosis and treatment efficacy.

A highlight of this study revolves around the predictive potential of these biomarkers. For instance, the differentiating capacity of nbTMBp in determining immunotherapy response underscores the importance of decomposing mutation burden into multiple features, such as non-B DNA. Similarly, the insights derived from the ovarian cancer case, where an increasing nbTMBp revealed heightened cisplatin sensitivity, highlights the potential treatment associated response. On the other hand, non-B-specific structures associated with mlTNB presents prediction capability in patient outcome in early-stage pancreatic cancer.

This research is not without limitations. While we demonstrated the patient grouping with clustering analyses, optimizing the thresholds for biomarkers with continuous value in clinical application remains a challenge. While the correlation between nbTMB/mlTNB and treatment responses is compelling, the mechanism remains unexplored. Future prospective studies are essential to elucidate the mechanisms underlying the observed associations as well as to validate the clinical applicability of these biomarkers.

In summary, we present two cancer biomarkers of non-B DNA associated burden that, with the role of non-B DNA in genomic instability, offers a novel expansion over existing markers for potentially explaining heterogenous outcomes and treatment responses in cancer.

## Data availability

The scripts for analyses in this study is available under a GPL-3.0 license in the Kowalski Lab GitHub repository (https://github.com/kmlabdms/MLNB).

## Acknowledgements

This work was supported by grant [RR160093 to support J.K.], research funds from the Department of Oncology, Dell Medical School [to J.K.].

## Author contributions

**JK** conceived of the idea, directed the analyses plan, and edited the manuscript.

**QX** performed the analyses, drafted the figures and initial manuscript.

